# Transient diquat inclusion and carboxylate-rim binding to carboxymethyl-*α*-cyclodextrin: a molecular dynamics study

**DOI:** 10.64898/2026.10.28.762698

**Authors:** Jing Chen, Xu Wu, Yongjian Huang, Wei Fu

**Affiliations:** Shenzhen UniBindMD Computation Biotech Co., Ltd., Shenzhen, China

**Keywords:** cyclodextrin, diquat, polyrotaxane, molecular dynamics, host–guest chemistry, pH-responsive materials, supramolecular detoxification

## Abstract

Rotaxane-based supramolecular scavengers have shown promise for diquat detoxification, but the relative contributions of cavity inclusion and interactions at the macrocycle rim remain unclear. We used all-atom molecular dynamics simulations in explicit water to examine diquat binding to carboxymethyl-*α*-cyclodextrin (CM-*α*-CD) and the behaviour of CM-*α*-CD on a poly(ethylene glycol) axle. Five systems were simulated: host–guest mixtures containing a neutral host (pH 3 model) or a hexaanionic host (pH 10 model), an explicitly constructed inclusion complex, and two-macrocycle pseudopolyrotaxanes initiated from pre-threaded or random configurations. The total production sampling was approximately 4.5 *µ*s.

A relaxed threaded geometry was obtained, with the diquat centroid 0.03 nm from the cavity axis and 9 of its 14 heavy atoms within the defined cavity cylinder. The guest left this geometry within the first 50 ps sampling interval and subsequently remained associated mainly with the carboxylate rim. During 1.2 *µ*s of unbiased sampling of the hexaanionic host–guest system, only 5 of 802 guest–frame pairs contained a guest atom within the cavity cylinder, and all of these events occurred near a portal. Rim contacts were more frequent: each guest formed an average of 3.8–4.4 host heavy-atom contacts per frame, with 12.8– 14.0% of guest heavy atoms in contact with the host. These interactions were substantially reduced for the neutral host. Sodium ions remained predominantly solvent-separated from the carboxylates. In the pseudopolyrotaxane simulations, pre-threaded rings satisfied the threading criterion in 1.6–8.4% of frames, and no spontaneous threading was observed from random configurations during 1 *µ*s. Under the simulated conditions, CM-*α*-CD therefore bound diquat primarily through dynamic rim interactions rather than persistent cavity inclusion.

## 1 Introduction

Diquat (1,1^*′*^-ethylene-2,2^*′*^-bipyridinium) is a fast-acting contact herbicide whose accidental or intentional ingestion can cause multiorgan failure, and no specific antidote is available. Engineered supramolecular scavengers that sequester the dication are therefore of interest, and rotaxane-based nanoscavengers have recently been reported to promote diquat detoxification in vivo [1]. The performance of such constructs depends partly on how the macrocycles interact with diquat. For modified cyclodextrins, association may involve encapsulation within the cavity or binding at a substituted rim, and these modes can respond differently to protonation and hydration. Distinguishing between them is necessary to relate macrocycle structure to scavenger function.

Cyclodextrins are cyclic oligosaccharides of *α*-1,4-linked glucopyranose units with a hydrophilic exterior and a comparatively apolar cavity, and they form inclusion complexes with a wide range of guests in water [2, 3]. Their cavity size is determined by the number of glucose units: *α*-, *β*- and *γ*-cyclodextrin have approximate inner diameters of 0.5, 0.6 and 0.8 nm, respectively. Guests must also pass through either the narrower primary portal or the wider secondary portal. Carboxymethylation of the primary rim introduces pH-titratable groups, allowing the macrocycle charge and its electrostatic interactions to vary with protonation state.

Cyclodextrins are also used to construct mechanically interlocked assemblies. When *α*-cyclodextrin rings are threaded onto a polymer chain, the resulting polyrotaxane contains macro-cycles that can slide along the axle but cannot escape if suitable end groups are present [4, 5]. This architecture underlies slide-ring materials, in which ring mobility contributes to elasticity [6], and mechanical bonding can also alter chemical reactivity, photophysics and guest affinity [7]. In rotaxane-based scavengers, the axle provides a scaffold for multiple macrocycles, while the macrocycles supply potential binding sites [1].

Macrocycles such as cucurbiturils and pillararenes have been widely studied as hosts for charged guests [8, 9], but the relative contributions of inclusion and rim association remain incompletely understood for carboxymethylated cyclodextrins. In particular, the persistence of an initially threaded diquat–cyclodextrin complex and the behaviour of unstopped CM-*α*-CD/PEG assemblies in water require direct examination. Atomistic simulations can address these questions by resolving guest position, host contacts and counterion distributions over time.

In this study, we used all-atom molecular dynamics simulations in explicit water to investigate these processes. We simulated CM-*α*-CD with two diquat dications in neutral and hexaanionic protonation states, an explicitly constructed CM-*α*-CD–diquat inclusion complex, and a pseudopolyrotaxane containing two CM-*α*-CD macrocycles and a PEG10 axle. The pseudopolyrotaxane was initiated from both pre-threaded and random configurations, giving approximately 4.5 *µ*s of total production sampling. Cavity occupancy, dethreading, host–guest contacts, counterion distributions, hydrogen bonding and axle conformation were analysed to compare inclusion with rim association.

## 2 Computational methods

### 2.1 Force field and topologies

All simulations used the CHARMM36 all-atom carbohydrate force field and CGenFF [10–12]. The CM-*α*-CD macrocycle (six glucopyranose units and six carboxymethyl groups on the primary rim) was parameterised in two protonation states: host-moe (162 atoms, neutral COOH groups) and host-lackH (156 atoms, six COO^−^ groups). Diquat was described by a CGenFF-derived ligand topology (26 atoms, +2e), and PEG10 was modelled with the peg topology (73 atoms). Water was represented by the TIP3P model [13]. Both host variants had fixed protonation states; proton transfer was not sampled. The pH 3 and pH 10 labels therefore denote end-member protonation models rather than a simulated titration curve, and neither the carboxymethyl pKa nor intermediate protonation patterns were determined.

### 2.2 System composition

Table 1 summarises the composition and sampling of the five systems; all of them contain explicit water. Figure 1 shows each cell in full and with the solvent removed.

**Table 1.** Composition and production sampling of the simulated systems.

| System | Host | Guests | Ions | Water | Box / ensemble | Sampling |
| --- | --- | --- | --- | --- | --- | --- |
| pH 3 host-guest | 1 × neutral CM-α-CD | 2 diquat | 4 Cl<sup>-</sup> | 2104 | 4 × 4 × 4 nm, NPT | 2 × 100 ns |
| pH 10 host-guest | 1 × CM-α-CD<sup>6-</sup> | 2 diquat | 2 Na<sup>+</sup> | 2089 | 4 × 4 × 4 nm, NPT | 1.2 μs |
| Inclusion complex | 1 × CM-α-CD<sup>6-</sup> | 1 diquat (threaded) | 4 Na<sup>+</sup> | 2101 | 4 × 4 × 4 nm, NVT then NPT | 1 ns NVT + 50 ns NPT |
| Polyrotaxane, pre-threaded | 2 × CM-α-CD<sup>6-</sup> | PEG10 | 12 Na<sup>+</sup> | 6876 | 5.92 × 5.92 × 5.92 nm, NPT | 2.0 μs |
| Polyrotaxane, random | 2 × CM-α-CD<sup>6-</sup> | PEG10 | 12 Na<sup>+</sup> | 6876 | 5.92 × 5.92 × 5.92 nm, NPT | 1.0 μs |

**Figure 1.**
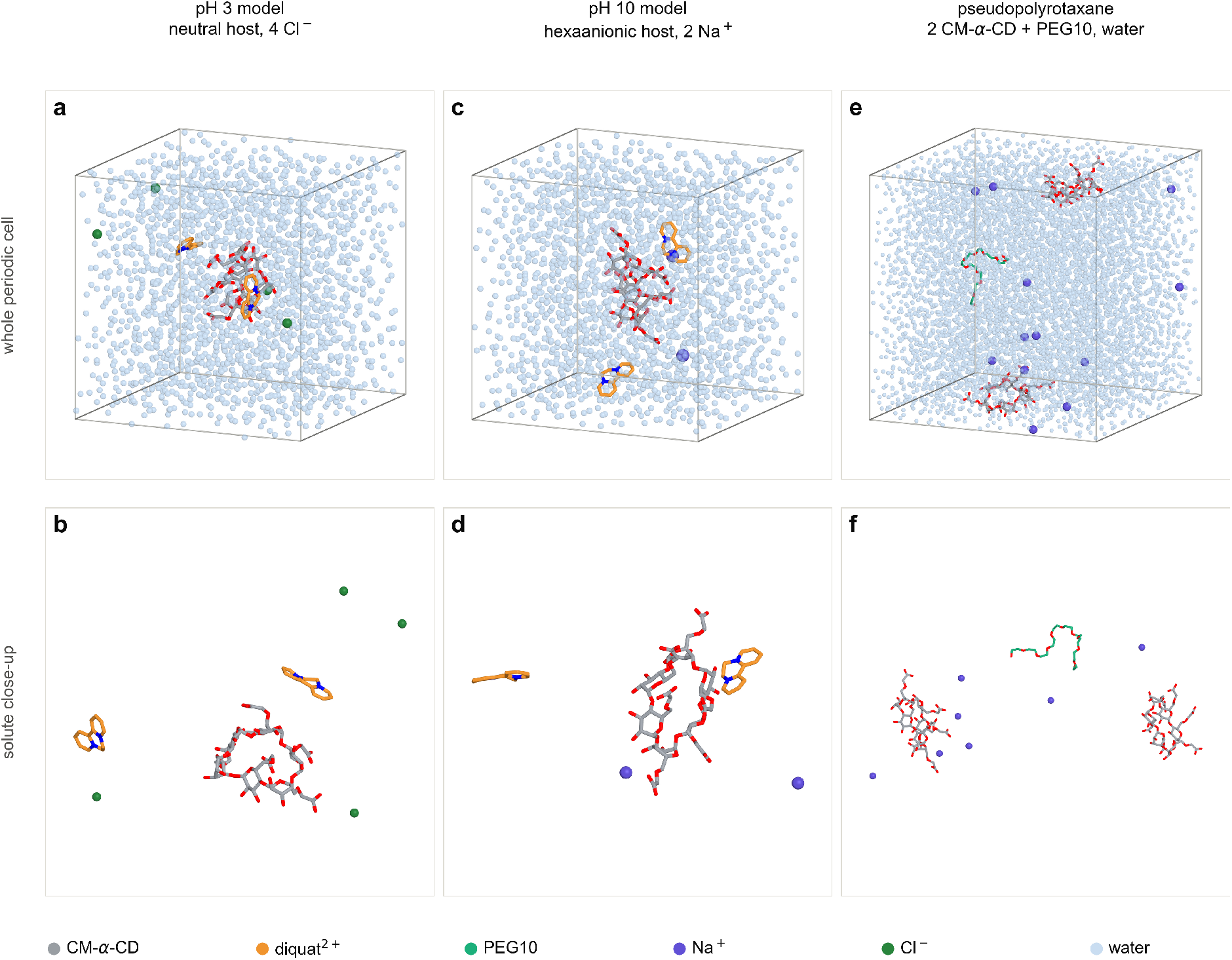
Model systems and simulation cells. Top row, whole periodic cell with water drawn as translucent oxygen spheres: (a) the pH 3 host–guest system (neutral host with two diquat dications and four Cl^−^); (c) the pH 10 host–guest system (hexaanionic host, two dications, two Na^+^); (e) the pseudopolyrotaxane (two CM-*α*-CD macrocycles and a PEG10 axle with twelve Na^+^). Bottom row, the same systems with water removed and the view opened up on the solute: (b) the neutral macrocycle with its two guests; (d) the hexaanionic macrocycle with both dications at the carboxylate rim; (f) the pseudopolyrotaxane after 2 *µ*s in water, in which the two macrocycles have separated and the PEG10 chain lies between them, unthreaded. The upper row is drawn at the scale of the simulation cell, the lower row is magnified; panels are not drawn to a common scale. Colours follow the key in the figure. Renderings were made with PyMOL.

Ion counts ensure electroneutrality in every box. In the pH 10 host–guest system the macro-cycle (−6) and two dications (+4) require two Na^+^; the inclusion complex carries one dication and therefore four Na^+^; in the polyrotaxane systems twelve carboxylates require twelve Na^+^.

The inclusion complex was constructed explicitly. One diquat dication was placed on the cavity axis of the hexaanionic host, with one pyridinium ring inside the cavity, the other protruding through the secondary rim, and the guest long axis parallel to the cavity normal. The assembly was solvated in a 4 nm cube with four Na^+^ ions, energy-minimised, and equilibrated for 1 ns at 300 K under constant volume. The equilibrated structure was then propagated for 50 ns at 300 K and 1 bar. This protocol allowed the persistence of an initially threaded geometry to be examined without requiring its spontaneous formation during the trajectory.

### 2.3 Simulation protocol

Simulations were run with GROMACS 2020.3 [14]. Electrostatics were treated with particle-mesh Ewald [15] (real-space cutoff 1.0 nm, 0.40 nm Fourier spacing and sixth-order interpolation), and Lennard-Jones interactions were treated with LJ-PME and a geometric combination rule [16], with analytical dispersion corrections for energy and pressure. Bonds to hydrogen were constrained with LINCS [17], allowing a 2 fs time step. Temperature was maintained at 300 K with a velocity-rescaling thermostat [18] (*τ* = 1.0 ps), and pressure was maintained at 1 bar with a Berendsen barostat [19] (*τ* = 2.5 ps, compressibility 4.5 *×* 10^−5^ bar^−1^). Centre-of-mass motion was removed every 100 steps. Coordinates were stored every 50 ps during production. Analyses used time series sampled at 50 ps for the inclusion complex, 2.5 ns for the 1.2 *µ*s alkaline run and 4 ns for the polyrotaxane runs.

For the pH 3 system, an initial trajectory generated without temperature control was excluded. The system was re-equilibrated for 1 ns at 300 K from a fresh velocity seed, followed by two independent 100 ns production replicas. All other trajectories were analysed after confirming that temperature, density and potential energy had become stationary (Figure 3).

### 2.4 Analysis definitions

The macrocycle frame was defined by the centroid and least-squares plane of the six glycosidic oxygens, with the normal oriented towards the primary (carboxymethylated) rim. Guest and axle atoms were expressed in cylindrical coordinates (r, z) about this frame. A guest atom was counted as inside the cavity if r < 0.45 nm and |z| < 0.45 nm. This cylinder encloses the *α*-CD cavity and the belt of ring oxygens and therefore provides a generous operational definition of inclusion. The threading coordinate for the inclusion complex was the radial distance *r*_*c*_ of the guest centroid from the macrocycle axis. Heavy-atom contacts used a 0.40 nm cutoff, and the guest contact fraction was defined as the fraction of guest heavy atoms making at least one such contact. Host atoms were grouped as carboxylate O, carboxyl C, glycosidic O, hydroxyl O or carbon skeleton. An axle was scored as threaded through a macrocycle when at least one axle atom lay within 0.50 nm of the axis on each side of the macrocycle plane (z > 0.15 nm and z < −0.15 nm). Hydrogen bonds required H*···* A < 0.25 nm and a D–H*···* A angle above 150°. Radial distribution functions were computed using the minimum-image convention and normalised by the ideal-gas shell volume. Solute-only trajectory frames were made whole and unwrapped molecule by molecule before frame-based analysis to avoid artefacts from periodic boundary conditions. Molecular images were prepared with in-house rendering code and cross-checked against VMD [20]; analyses were implemented in NumPy [21] and rendered with Matplotlib [22].

## 3 Results and discussion

### 3.1 Equilibration and stability of the trajectories

Temperature, density and potential energy became stationary within the first few nanoseconds and remained stable during the production windows (Figure 3). Mean values over the second half of each run were 299.9 *±* 3.7 K and 1025.5 *±* 2.6 kg m^−3^ for the pH 10 host–guest system, 299.9 *±* 3.6 K and 1025.2 *±* 2.7 kg m^−3^ for the inclusion complex, and 300.0 *±* 2.0 K and 1018.3 *±* 1.4 kg m^−3^ for the aqueous polyrotaxane. For the two pH 3 replicas, the corresponding values were 300.1 *±* 3.5 K and 1021.2 *±* 2.5 kg m^−3^ (replica A) and 299.8 *±* 3.9 K and 1021.3 *±* 2.6 kg m^−3^ (replica B). Instantaneous pressures fluctuated about zero with amplitudes similar to those in the pH 10 run.

The two pH 3 replicas were used to assess run-to-run variation in the guest descriptors. Contact counts, contact fractions and cavity occupancy were calculated per frame and pooled where indicated; replica-specific values are reported where relevant.

### 3.2 Geometry and rarity of diquat inclusion

After energy minimisation and 1 ns of equilibration at 300 K, the constructed complex retained a threaded geometry. The guest centroid was 0.03 nm from the macrocycle axis, the N6–N8 axis of diquat was inclined by 20° to the cavity normal, and 9 of the 14 guest heavy atoms fell inside the cavity cylinder (r < 0.45 nm, |z| < 0.45 nm; Figure 2c). One pyridinium ring lay inside the cavity (z = +0.14 nm, where z > 0 points towards the carboxymethylated rim), while the opposite end protruded beyond the secondary rim (z = −0.55 nm). The guest therefore passed through the macrocycle rather than resting against one face.

**Figure 2.**
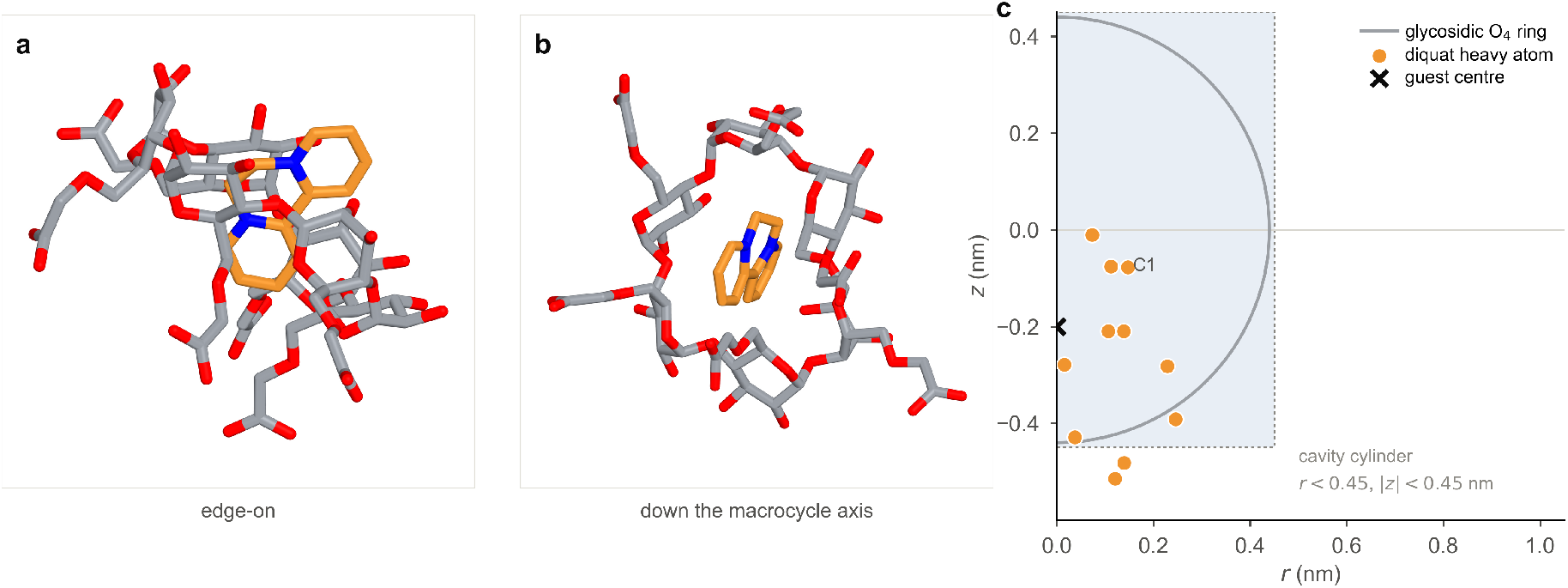
The threaded inclusion complex. (a) Edge-on and (b) down the macrocycle axis views of the relaxed CM-*α*-CD–diquat complex: the dication lies along the cavity axis, with one pyridinium ring inside the cavity and the other protruding through the secondary rim. (c) Positions of the 14 diquat heavy atoms in the macrocycle frame after energy minimisation and equilibration, plotted as radius r against axial coordinate z; the shaded box is the cavity cylinder (r < 0.45 nm, |z| < 0.45 nm), the grey arc is the ring of glycosidic O_4_ oxygens, and the cross marks the guest centre. Nine of the 14 heavy atoms lie inside the cylinder, and the guest axis makes an angle of 20° with the macrocycle normal. Water, ions and hydrogen atoms are omitted.

**Figure 3.**
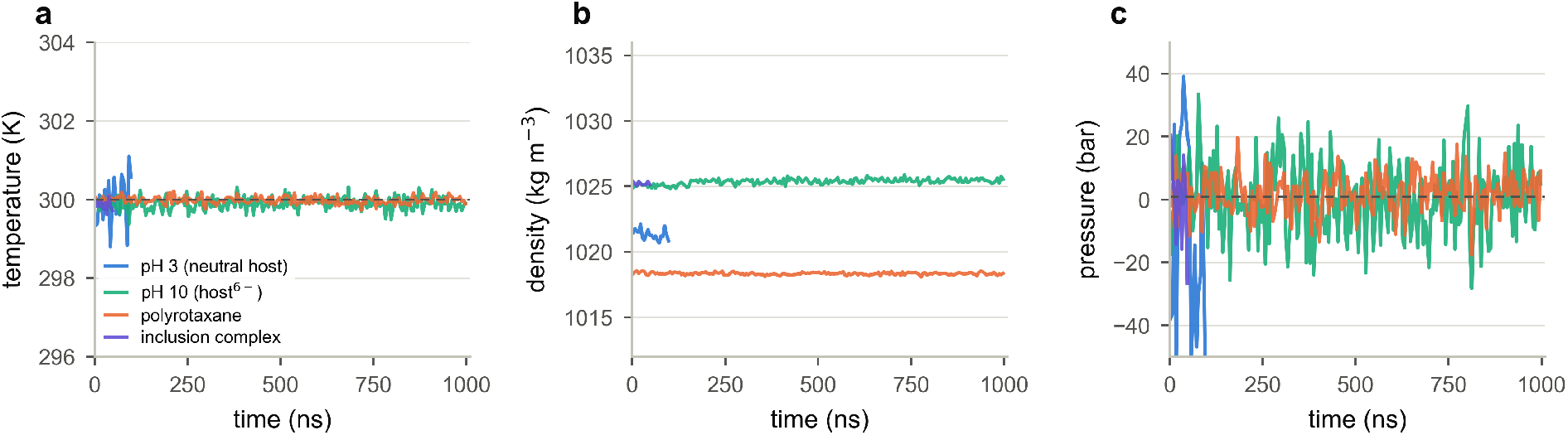
Equilibration of the production runs. (a) Temperature, (b) density and (c) pressure, averaged in 5 ns blocks, for the pH 3 host–guest system (2 *×* 100 ns, replica A shown), the pH 10 host–guest system (1.2 *µ*s), the aqueous pseudopolyrotaxane (2 *µ*s) and the inclusion complex (50 ns, visible only near the origin of the 1 *µ*s axes). Dashed lines mark 300 K and 1 bar. All four systems are aqueous; no vacuum trajectory is included.

This geometry is sterically plausible. Although the approximately 0.5 nm *α*-CD portal is narrower than the roughly 0.7 nm long axis of diquat, it can accommodate the cross-section of a pyridinium ring (approximately 0.45 nm), allowing end-on insertion without substantial ring distortion. The relaxed structure positioned the charged nitrogens near ring oxygens and placed part of the bipyridinium core within the cavity. However, the stability of this constructed geometry during minimisation and equilibration does not by itself establish its persistence or its stability relative to rim-associated states.

Cavity entry was rare in the unbiased simulations. Over 1.2 *µ*s at pH 10, the guest density in the macrocycle frame (Figure 4a) was concentrated in an annulus at r = 0.5–1.1 nm, corresponding to the carboxylate rim and surrounding solvent shell. Only 0.09% of guest heavy-atom density lay inside the cavity cylinder, and 5 of 802 guest–frame pairs contained at least one atom in this region (one for guest 1 and four for guest 2, with at most two atoms at a time). All five events occurred near a portal (|z| = 0.34–0.71 nm, r = 0.30–0.77 nm). The pH 3 replicas produced six such atom–frames in 4004 guest–frame pairs (0.15%). These observations indicate transient contacts at the cavity entrances but do not demonstrate spontaneous formation of a fully threaded complex in the unbiased trajectories.

**Figure 4.**
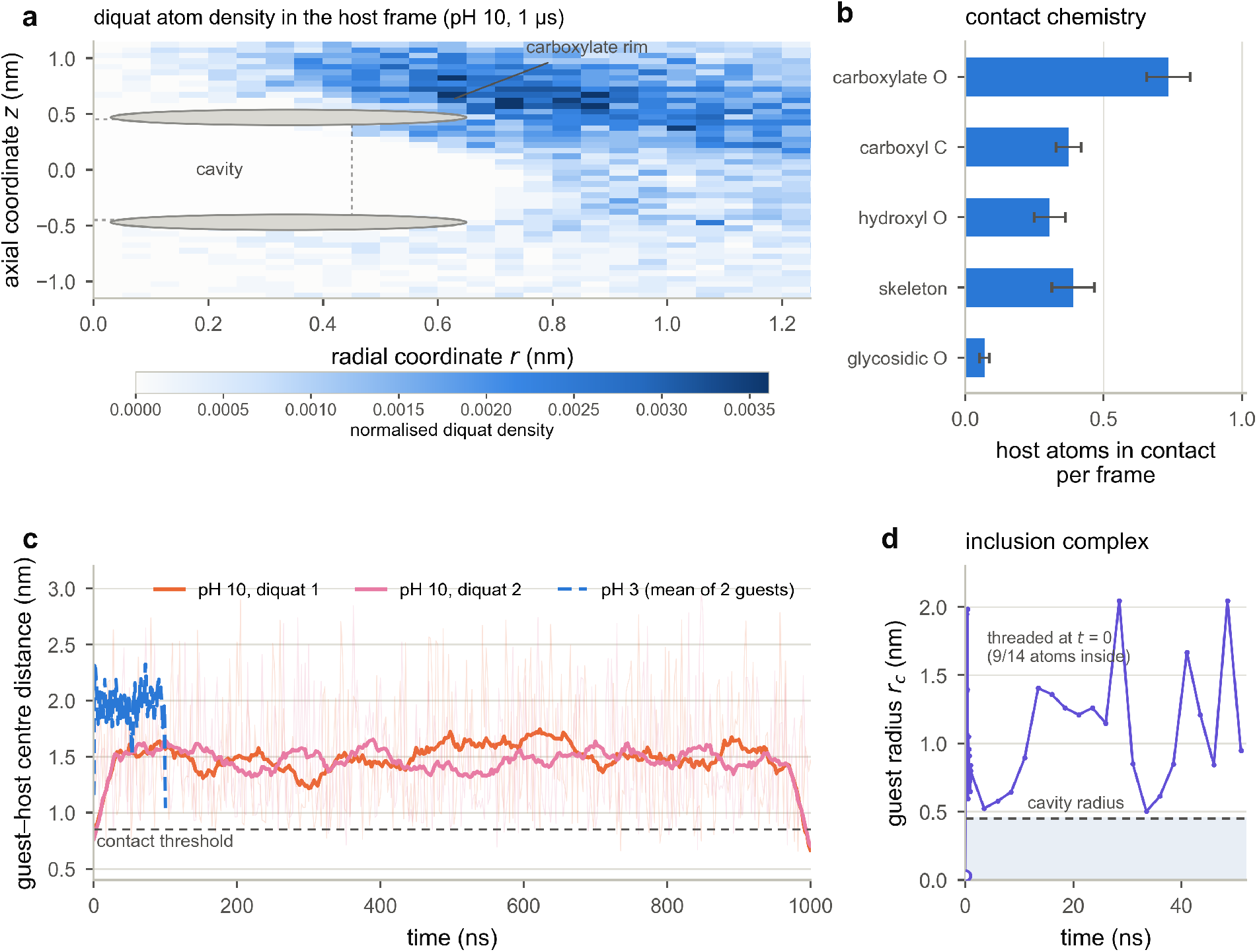
Cavity entry, rim capture and dethreading. (a) Normalised density of diquat heavy atoms in the macrocycle frame at pH 10 (1.2 *µ*s), projected onto the axial (z) and radial (r) coordinates; the dashed box marks the cavity region (r < 0.45 nm, |z| < 0.45 nm), which holds 0.09% of the guest atom density, and the grey ellipses are a schematic cross-section of the macrocycle. (b) Contact chemistry: mean number of host atoms of each class within 0.40 nm of a guest, with 95% confidence intervals of the mean. (c) Centre-to-centre distance between each guest and the macrocycle over 1 *µ*s; thin lines are raw data (2.5 ns resolution), thick lines are running means, and the dashed blue trace is the mean of the two guests in the pH 3 system. (d) Radial distance of the guest centroid from the macrocycle axis for the threaded inclusion complex: 1 ns NVT followed by 50 ns NPT, at 50 ps resolution; the dashed line marks the cavity radius, the shaded band the cavity interior, and the hollow marker the relaxed threaded structure at t = 0, in which 9 of 14 heavy atoms are inside the cavity.

### 3.3 The constructed inclusion complex rapidly relaxes to rim-associated states

The threaded geometry did not persist during production. In the trajectory started from the relaxed inclusion complex, the guest left the cavity within the first 50 ps sampling interval: *r*_*c*_ increased from 0.03 to 0.63 nm, and no guest atom remained inside the cylinder (Figure 4d). The guest nevertheless remained associated with the macrocycle for the following 50 ns, with *r*_*c*_ fluctuating between 0.5 and 2.0 nm (mean 1.08 nm). The closest heavy-atom approach reached 0.28 nm, and one transient re-entry into the cavity cylinder occurred at 35 ns. The dethreading trajectory thus ended in rim-associated configurations similar to those sampled in the unbiased pH 10 simulation.

The dethreading and unbiased simulations therefore sampled similar rim-associated configurations, despite starting from different guest positions. Because inclusion was imposed in the initial structure and only one dethreading trajectory was obtained, these data do not provide a free-energy difference or a statistically converged lifetime. They show that the constructed threaded geometry was not persistent on the nanosecond timescale under the simulated conditions; the 50 ps frame interval provides only an upper bound on the time at which the first saved non-threaded configuration was observed.

A possible contribution to this behaviour is the mismatch between the strongly hydrated dication and the comparatively apolar cavity. Cavity inclusion may require partial desolvation of the pyridinium groups, whereas association with the carboxylate rim can provide electrostatic interactions while leaving much of the guest exposed to water. The present trajectories are consistent with this interpretation but do not resolve the underlying dehydration or binding free energies.

### 3.4 pH-dependent rim association

Rim association was dynamic. In the pH 10 run, the guest–host centre-to-centre distance fluctuated around 1.5 nm, with excursions below 1.0 nm in 17–19% of frames and minimum instantaneous heavy-atom separations of 0.26–0.27 nm (Figure 4c). Guests 1 and 2 formed averages of 4.4 and 3.8 host heavy-atom contacts per frame within 0.40 nm, respectively, and 14.0% and 12.8% of their heavy atoms contacted the host. Multiple transitions between associated and more distant configurations occurred during the microsecond trajectory, consistent with transient association rather than a single long-lived bound state.

The contact decomposition indicates that the carboxylate rim was the main interaction site (Figure 4b). Carboxylate oxygens formed the largest single class of contacts (0.73 host atoms per frame), followed by the carbon skeleton (0.39), carboxyl carbons (0.37) and hydroxyl oxygens (0.31); glycosidic oxygens contributed 0.07. The average angle between the N6–N8 axis and the macrocycle normal was 57°, consistent with an oblique orientation across the rim. This arrangement allows interactions with the carboxylate groups while retaining substantial solvent exposure of the guest.

Protonation markedly reduced these interactions. In the neutral-host replicas, the two dications were located 1.96 *±* 0.49 and 1.99 *±* 0.50 nm from the macrocycle centre (replica A) and 2.00 *±* 0.50 and 1.95 *±* 0.50 nm (replica B), approximately 0.4–0.5 nm farther from the host than at pH 10. Residual contacts were sparse: 0.52–0.72 host atoms per frame, compared with 3.8–4.4 at pH 10, and the guest contact fraction was 2.2–3.2%, compared with 12.8–14.0%. The carboxyl-group contribution decreased from 0.73 to 0.07 host atoms per frame, while hydroxyl oxygens (0.11) and the carbon skeleton (0.07) accounted for most of the remaining contacts. Chloride remained dissociated, with a contact within 0.30 nm in at most 0.1% of frames. These changes are consistent with weaker hydrogen-bonding and dispersion interactions in the neutral-host model and a substantial electrostatic contribution in the hexaanionic model.

### 3.5 Sodium counterions are predominantly solvent-separated

Because the binding is electrostatic, the fate of the counterions is central. In water the sodium ions are not condensed onto the carboxylate rim. In the pH 10 host–guest system only 2.2% of frames contain a Na^+^ within 0.30 nm of a carboxylate oxygen, and the median closest distance is 0.80 nm (Figure 5b); in the polyrotaxane box, which holds twelve Na^+^ for twelve carboxylates, the median falls to 0.42 nm but still sits at the outer-sphere separation. The Na^+^–O radial distribution function (Figure 5a) shows the contact region at 0.20–0.30 nm as a narrow, weakly populated peak and a dominant first maximum at 0.40–0.45 nm with *g*(*r*) ≈ 6–7, characteristic of a solvent-separated (outer-sphere) ion pair in which a single water molecule sits between the ion and the carboxylate. The trajectory of the closest distance (Figure 5c) shows the ion spending most of its time at 0.8–1.0 nm, with brief excursions to contact that last only a few frames.

**Figure 5.**
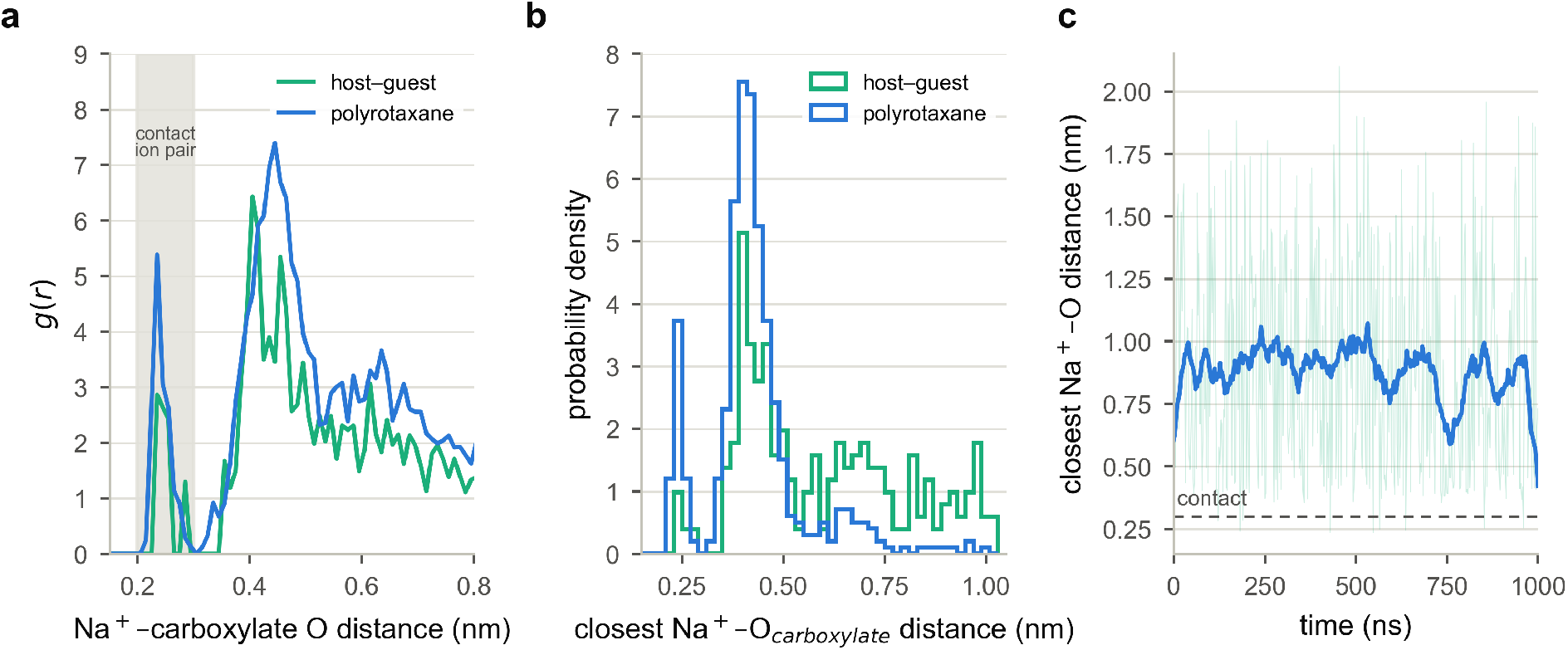
Sodium counterions in water. (a) Na^+^–carboxylate oxygen radial distribution function for the host–guest (pH 10) and polyrotaxane systems; the shaded band marks the contact-pair region, which is weakly populated. (b) Probability density of the closest Na^+^–O_carboxylate_ distance per frame. (c) Closest Na^+^–O distance versus time for the 1 *µ*s pH 10 run; the thin trace is raw data at 2.5 ns resolution, the thick line a running mean, and the dashed line the 0.30 nm contact threshold.

Sodium therefore did not form a persistent, site-bound layer that neutralised the carboxylate rim in these simulations. Electrostatic screening instead involved hydration and a diffuse, mobile counterion distribution. Explicitly accounting for solvent and counterion organisation is consequently important when interpreting guest binding to the carboxymethylated host.

Hydrogen bonding within the pseudopolyrotaxane did not alter this picture. Host–axle hydrogen bonds remained below one per frame even in the pre-threaded simulation and were essentially absent after the rings left the axle (<0.01 per frame). Mechanical confinement and dispersion, rather than an extensive hydrogen-bond network, were therefore the main host–axle interactions observed.

### 3.6 Limited persistence of CM-*α*-CD threading on PEG10

The pseudopolyrotaxane simulations examined the behaviour of the same macrocycles on a PEG10 axle (Figure 6). When two CM-*α*-CD rings were initially threaded onto the axle, the threading criterion was satisfied in only 1.6% (host 1) and 8.4% (host 2) of frames over 2 *µ*s. The macrocycles separated to a mean centre-to-centre distance of 3.34 *±* 0.62 nm, and PEG10 adopted dimensions close to those of a free chain (*R*_*g*_ = 0.63 *±* 0.09 nm; *R*_*e*_ = 1.54 *±* 0.49 nm). Host–axle contacts decreased to averages of 0.95 and 4.2 heavy atoms per frame. Threading events were brief and clustered: the closest macrocycle–PEG distance fell below 0.5 nm for tens of nanoseconds around 1 *µ*s before returning to approximately 1.3 nm (Figure 6b).

**Figure 6.**
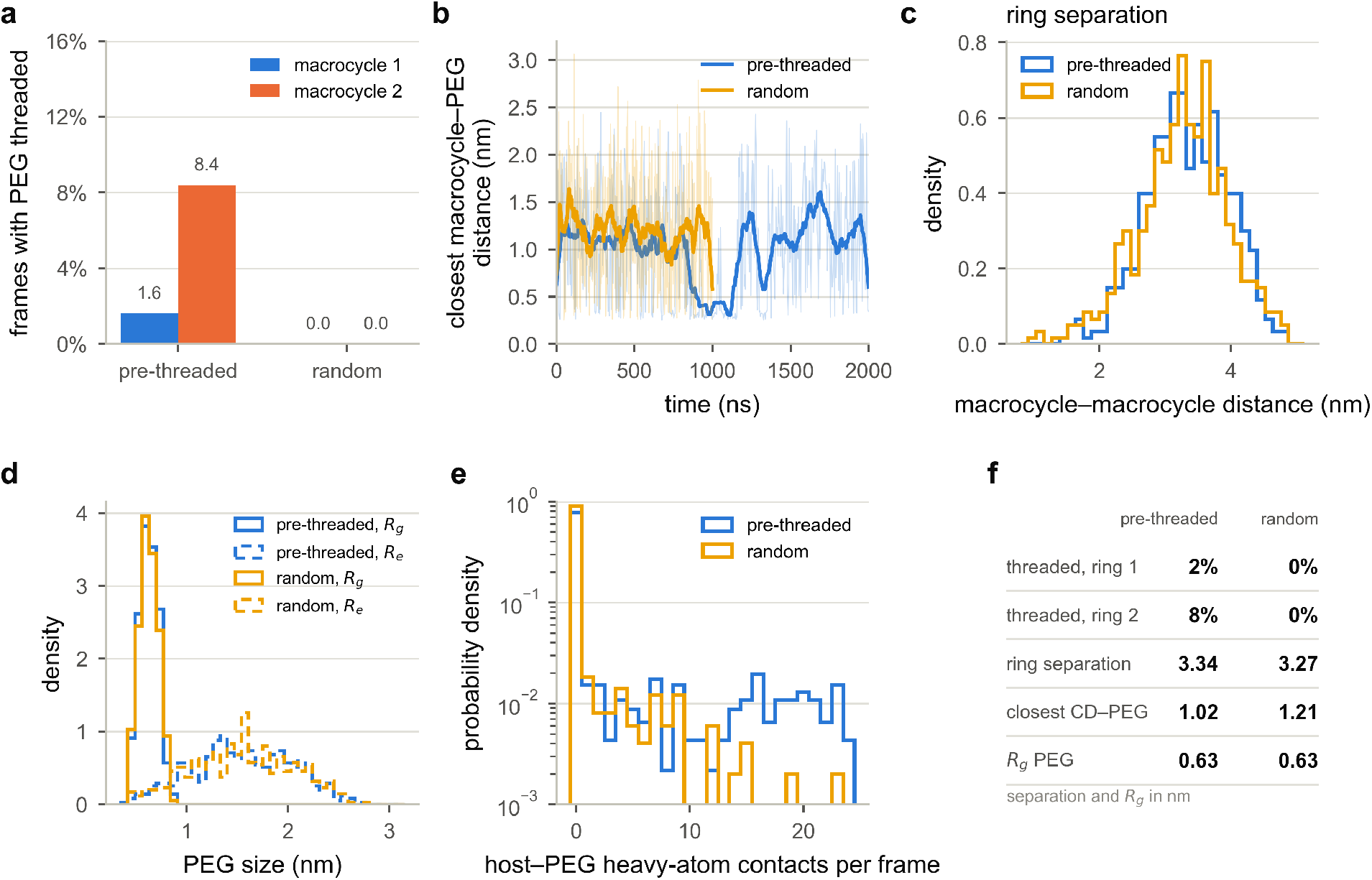
Threading of CM-*α*-CD on PEG10 in water. (a) Fraction of frames in which the axle is threaded through each macrocycle for pre-threaded and random-initiated simulations. (b) Closest macrocycle–PEG distance versus time (thin: raw, thick: running mean). (c) Distribution of macrocycle– macrocycle centroid distances. (d) Distribution of PEG radius of gyration (solid) and end-to-end distance (dashed). (e) Host–PEG heavy-atom contacts per frame on a logarithmic scale. (f) Summary of the mean values.

No threading event was observed during the 1 *µ*s simulation started from randomly placed macrocycles. The two hosts remained separated by an average of 3.27 nm, the axle radius of gyration was 0.63 nm, and host–axle contacts remained low. Spontaneous threading was therefore not observed on the microsecond timescale under these conditions. This result is consistent with the use of templating or controlled reaction conditions in polyrotaxane synthesis [4, 5], although the simulation does not sample synthetic assembly pathways directly.

These percentages describe an unstopped pseudopolyrotaxane and should not be extrapolated to end-capped rotaxanes. Without bulky end groups, rings can slide off the axle, so ring loss is an expected possible outcome rather than, by itself, evidence for or against the force field. The contrast with the diquat inclusion complex also reflects different assembly processes: a small charged guest can be positioned within the cavity, whereas spontaneous threading of a flexible neutral polymer was not observed at the concentration and timescale examined.

### 3.7 Implications for scavenger design

The simulations suggest that rim association may contribute more than persistent cavity inclusion to diquat binding by CM-*α*-CD under the conditions examined. A rotaxane scaffold could present several carboxymethylated rims on one construct, potentially increasing the local density of interaction sites. End groups would nevertheless be required to prevent macrocycle loss from the axle.

These observations suggest several directions for subsequent design studies. If persistent inclusion is required, alternative cavity sizes or chemical modifications could be tested, although the present simulations do not compare host sizes and cannot predict which modification would improve binding. Optimising rim charge density and hydration may be more directly relevant to the binding mode observed here. The pseudopolyrotaxane results also support the inclusion of stoppers in any design that relies on retained rings. Finally, counterion binding and solvent organisation should be examined explicitly rather than inferred solely from the nominal host charge.

## 4 Conclusions

All-atom molecular dynamics simulations showed that a threaded CM-*α*-CD–diquat geometry could be constructed and maintained through minimisation and equilibration, but it did not persist during the subsequent production simulation. The guest left the cavity within the first 50 ps sampling interval and adopted rim-associated configurations. In unbiased simulations of the hexaanionic host, cavity contacts were rare and restricted to portal regions, whereas rim contacts were frequent and dynamic. The neutral host showed markedly fewer guest contacts, supporting an important contribution from carboxylate–diquat electrostatic interactions.

Sodium counterions were predominantly solvent-separated, indicating that persistent site-bound sodium did not neutralise the carboxylate rim in these simulations. CM-*α*-CD threading on an unstopped PEG10 axle was also weakly persistent, and no spontaneous threading was observed from random configurations within 1 *µ*s. Under the conditions examined, diquat association with CM-*α*-CD was therefore dominated by transient rim interactions rather than stable cavity encapsulation. Free-energy calculations, additional replicas and end-capped constructs will be needed to quantify binding thermodynamics, dethreading kinetics and the behaviour of complete rotaxane scavengers.

## Acknowledgments

This work was supported by Shenzhen UniBindMD Computation Biotech Co., Ltd. Computational resources, including GPUs and access to a high-performance computing cluster, were provided by the UniBindMD supercomputing platform.

## Author contributions

Jing Chen: conceptualization, methodology, software, investigation, formal analysis, writing— original draft. Xu Wu: methodology, validation, data curation, writing—review and editing. Yongjian Huang: supervision, resources, project administration, funding acquisition, writing— review and editing. Wei Fu: conceptualization, supervision, writing—review and editing.

## Data availability

Trajectory-derived descriptor tables, analysis scripts (Python/NumPy), and the parameters used to generate every figure in this manuscript are available from the corresponding author on reasonable request. Raw trajectories (XTC) are archived on the Beijing Supercomputing Center system and can be transferred on request.

## Supporting information

Additional descriptor time series for the host–guest segments, per-replica pH 3 statistics, and the complete contact-decomposition tables are provided as Supporting Information.

